# First record of Zaprionus tuberculatus (Diptera: Drosophilidae) in mainland France

**DOI:** 10.1101/2023.10.09.561531

**Authors:** Romain Georges, Amir Yassin, Hervé Colinet

## Abstract

**BACKGROUND:** As most drosophilid species are not considered as primary pest, studies of the whole drosophilid communities, including other genera than *Drosophila*, with standardized surveys are relatively sparse. However, the spotted wing drosophila *Drosophila suzukii* (Matsumura, 1931) and its rapid expansion through the world led to the implementation of many monitoring programs in various countries. As part of a research project on *D. suzukii*, we set up in 2022 an annual fly monitoring in 16 fruits farms to understand populations dynamics of *D. suzukii* and to survey drosophilid communities.

**RESULTS:** We report here the first observation of *Zaprionus tuberculatus* Malloch, 1932 (Diptera: Drosophilidae) in mainland France. Over the whole monitoring, we trapped a total of 111 specimens in a fig orchard located in southern France (Salses-le-Château), both in fig trees and nearby hedgerows. The first detection of *Z. tuberculatus* occurred in July 2022 in the hedgerow and captures continued until January 2023 with an interruption in November and December. In addition, in this orchard we collected overripe figs in September 2023 from which over 15 *Z. tuberculatus* have emerged in the following two weeks.

**CONCLUSION:** The pest status of *Z. tuberculatus* and its potential risk for agriculture is not clear, but the pest behavior of the close-relative species *Zaprionus indianus*, especially on figs, should be a warning point for the entry of *Z. tuberculatus* into the EU and France, as they may have similar nutritional ecology. The pest status, the establishment and the future spread of *Z. tuberculatus* should thus be monitored to assess possible damages to fruits productions.

## 1. Introduction

Drosophilidae Rondani is a very diverse family with almost 4,700 described species distributed in 77 genera (1). The largest genus of Drosophilidae is *Drosophila* Fallén including 1,676 described species. There is a strong heterogeneity in the existing knowledge on Drosophilidae, with a huge literature on genetics as well as cellular and developmental biology of *Drosophila* species, mainly *Drosophila melanogaster*, but very few knowledges on other genera. Studies of the whole drosophilid communities, including other genera than *Drosophila*, with standardized surveys are relatively sparse (2–4). A possible reason for this lack of interest is likely the ecology and lifecycle of Drosophilidae. For most drosophilids species, larvae grow on decaying plant material and forage on associated microbial communities (5,6) and are therefore not considered as primary pest. In addition, these flies are small and cause no trouble to humans or animals, so they go unnoticed by most people. However, the spotted wing drosophila *Drosophila suzukii* (Matsumura, 1931) and its rapid expansion through the world changed the game. The damage caused to cultivated fruits by this invasive species and the huge economic loss have led to a large number of studies in the last decade about its ecology and the control measures (7). Many monitoring programs have been set up in various countries to better understand and predict populations dynamics of *D. suzukii* (8–11). As part of a research project on *D. suzukii* (DroFramb action within the framework of ANR Drothermal project, https://www.drothermal.cnrs.fr/), in 2022, we set up an annual fly monitoring in various regions of France. The main objective was to conduct a vast monitoring program of *D. suzukii* during a whole year, including during winter months, to get data on fly presence in various areas and climates. This monitoring effort also had the secondary objective of surveying *Drosophilidae* communities. Field monitoring is crucial to detect novel invaders and potential novel pests. Indeed, early detection is key to successful preventative strategies. In the present study, we will not show the data on *D. suzukii* over the course of this national monitoring program, but instead, we provide a first record of the establishment of a new invasive and potential pest fly, *Zaprionus tuberculatus* Malloch, 1932 (Diptera: Drosophilidae) in mainland France.

## 2. Materials and Methods

From March 2022 to February 2023, we conducted a national participative monitoring of *D. suzukii* (Matsumura, 1931) with 16 fruits producers along a latitudinal gradient in France. In each farm, three traps were installed at least 10m apart in an orchard (fig and cherry) or a raspberry plantation at fruiting height and three others in a hedgerow or a grove nearby at approximately 1 m height. Traps were opened for ten days at the end of each month resulting in twelve 10-days trapping sessions across seasons. Flies were captured using a red-colored water bottle trap pierced with six 5mm holes. The bottles contained a bait (80 ml) and drowning solution (30 ml) in a collection tube. For the bait, we used a mixture of 80% apple cider vinegar (5% acidity) and 20% of cane sugar syrup. Flies caught in the bottles fell in the drowning solution that was made of salted water (40g.L^-1^) with a drop of odorless wetting agent (L’Arbre Vert® - Peaux Sensibles sans parfum, France). Bait and drowning solution were renewed at each trapping session. Local temperatures at each site were monitored during all the experiment using TMS-4 Standard dataloggers (TOMST®, Czech Republic) which collected data every 15 minutes in 3 different levels: 6 cm into the soil, at the surface and 15 cm above the soil surface.

## 3. Results and discussion

While identifying drosophilid specimens collected in different regions of France, we found flies with white longitudinal stripes at a single location in a fig orchard in southern France (Salses-le-Château; 42°50’0.169”N 2°55’5.447”E ; see Figure 1 & 2). Flies with these morphological features typically belong to the *Zaprionus* Coquillet, 1902 genus. According to the Köppen-Geiger classification (12,13), the climate of this region is type Csa, corresponding to hot-summer Mediterranean climate with mild winters averaging above 0°C and hot summers averaging above 25°C. In this location, over the whole monitoring period, we trapped 111 specimens of *Zaprionus sp*. out of a total of 3766 drosophilids (Table 1).

**Table 1.** Occurrence of *Z. tuberculatus* collected in France. Number of male and female *Z. tuberculatus* and all other Drosophilidae trapped in a fig orchard and a hedgerow nearby during one-year monitoring in 2022-23. Temperatures indicated are average, minimum (min) and maximum (max) of average daily temperature recorded every 15 minutes by TMS-4 dataloggers (TOMST, Czech Republic) at 15cm above the soil surface.

| Date collected | Fig orchard |  |  |  | Hedgerow |  |  |  |
| --- | --- | --- | --- | --- | --- | --- | --- | --- |
|  | *Zaprionus tuberculatus* |  | Other Drosophilidae | Daily temperature (average ; min - max) | *Zaprionus tuberculatus* |  | Other Drosophilidae | Daily temperature (average ; min - max) |
|  | Males | Females |  |  | Males | Females |  |  |
| March 2022 | 0 | 0 | 36 | 11.8 ; 8.9 - 17.6 | 0 | 0 | 67 | 11.9 ; 8.9 - 17.6 °C |
| April 2022 | 0 | 0 | 58 | 15.1 ; 6.5 - 22.2 | 0 | 0 | 33 | 15.1 ; 5.9 - 22.3 °C |
| May 2022 | 0 | 0 | 1 | 20.9 ; 17.1 - 26.2 | 0 | 0 | 32 | 20.2 ; 17.3 - 25.6 °C |
| June 2022 | 0 | 0 | 131 | 25.5 ; 21.3 - 30.3 | 0 | 0 | 172 | 24.4 ; 20.2 - 29.3 °C |
| July 2022 | 0 | 0 | 650 | 28.7 ; 24.7 - 32.7 °C | 11 | 19 | 1356 | 27.5 ; 23.2 - 32.1 °C |
| August 2022 | 12 | 13 | 390 | 28.3 ; 24.0 - 33.4 °C | 13 | 14 | 479 | 27.7 ; 23.7 - 32.6 °C |
| September 2022 | 0 | 0 | 5 | 22.4 ; 16.2 - 27.7 °C | 4 | 4 | 168 | 22.4 ; 16.3 ; 27.7 °C |
| October 2022 | 4 | 2 | 227 | 19.5 ; 17.2 - 21.8 °C | 6 | 6 | 330 | 19.6 ; 17.3 - 21.9 °C |
| November 2022 | 0 | 0 | 2 | 13.0 ; 7.6 - 19.0 °C | 0 | 0 | 81 | 13.6 ; 8.6 - 19.9 °C |
| December 2022 | 0 | 0 | 0 | 9.2 ; 2.7 - 14.1 °C | 0 | 0 | 2 | 10.0 ; 4 - 15.3 °C |
| January 2023 | 0 | 0 | 27 | 7.6 ; 3.0 - 12.8 °C | 2 | 1 | 24 | 8.1 ; 3 - 13.8 °C |
| February 2023 | 0 | 0 | 71 | 8.9 ; 3.6 - 14.4 °C | 0 | 0 | 50 | 9.5 ; 3.5 - 15.2 °C |

**Figure 1.**
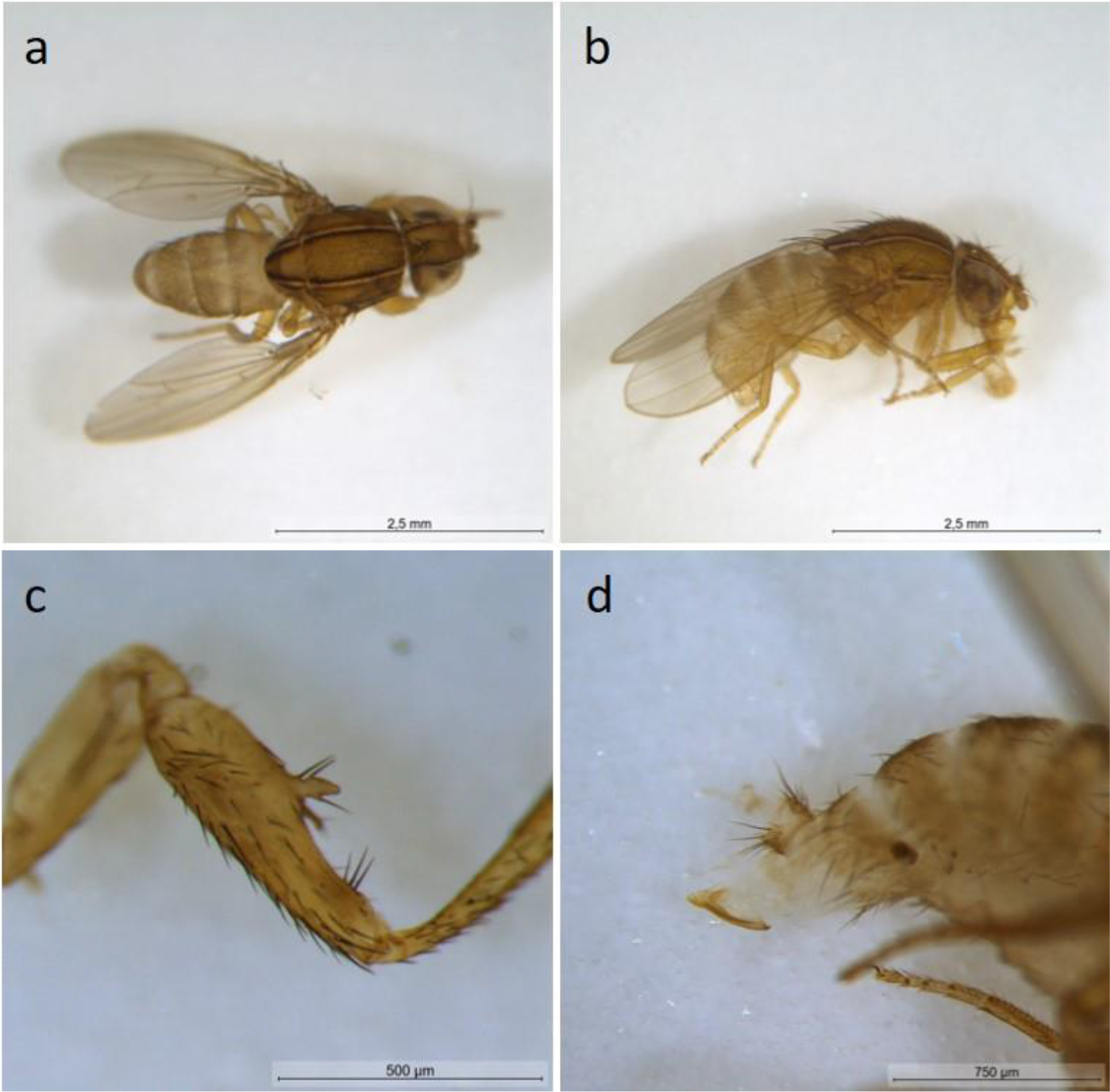
Z. tuberculatus: dorsal view (a), lateral view (b) and detail of the protruding tubercle on the forefemur of a male; detail of the ovipositor of a female.

**Fig 2.**
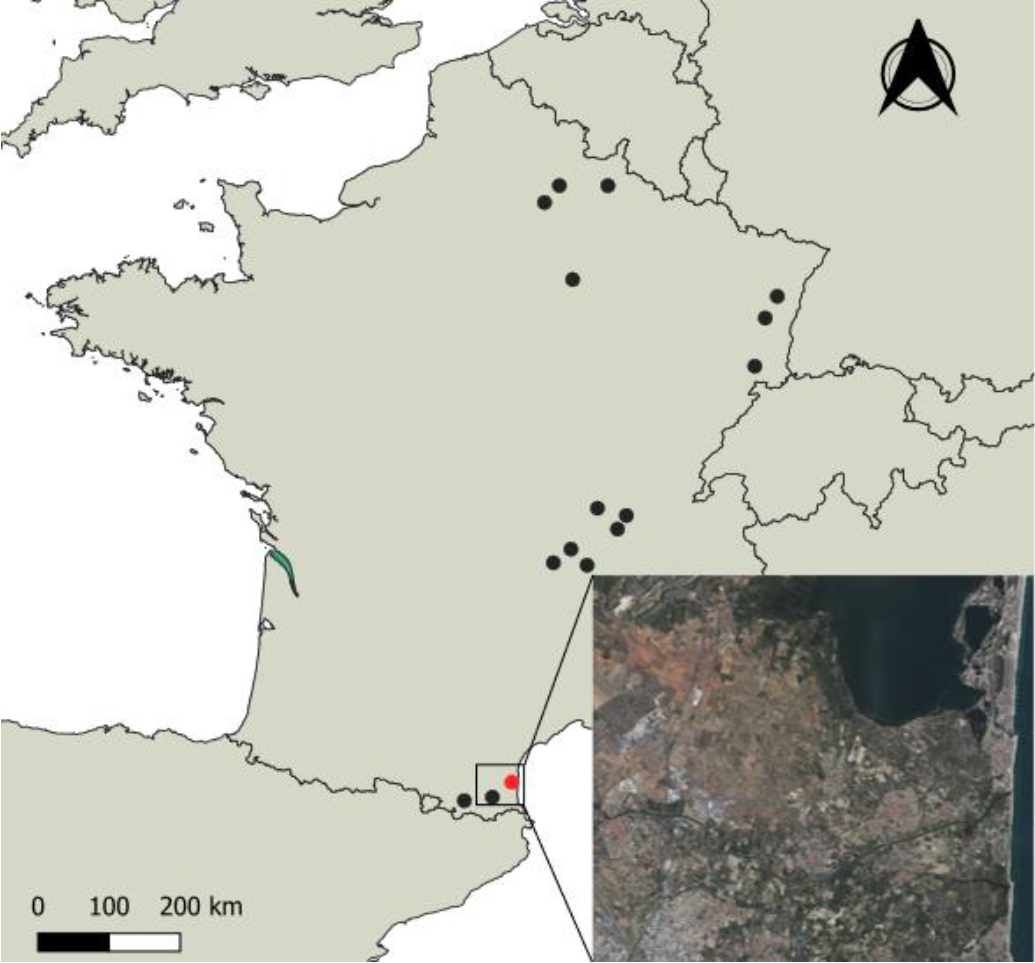
French locality (red dot) where specimens of Z. tuberculatus were collected. Black dots represent the other monitored localities where no Z. tuberculatus specimens were collected during the same period. (© EuroGeographics for the administrative boundaries and IGN BD-ORTHO for the satellite image)

Species identification was first done by R. Georges and then sent to A. Yassin for confirmation based on external morphological criteria. The genus *Zaprionus* Coquillett, 1902 exhibits characteristic white longitudinal stripes on the frons and the mesonotum (Fig. 1a, b) and we could also easily distinguish a protruding tubercle bearing a bristle on the forefemur of all our specimens (Fig. 1c). This feature is characteristic of a clade of Afrotropical species, the *Zaprionus tuberculatus* subgroup (14). The subgroup contains seven species, of which only one, *Zaprionus tuberculatus* (Malloch, 1932) has invasive capacities and has expanded its geographical range to the palearctic region during the last four decades (15). The seven species can be distinguished on the basis of morphological characters of male and female genitalia and internal reproductive organs as well as immature stages (14,16). Of these characters, male and female genitalia provide the best diagnoses on non-living specimens, with DNA barcoding being of limited utility due to shared mitochondrial sequences among closely-related species (15). Dissection of a sample of captured specimens from Salses-le-Château supported the morphological identification of the introduced flies as *Z. tuberculatus*. Female spermathecae (Figure 3a) have predominantly smooth surface except on the tip wherein the surface becomes slightly rigged, whereas the phallotrema of the male aedeagi (Figure 3b) has conspicuous ventral process and a densely teethed flap surrounding its border. Individuals are now kept in 70% alcohol and are available from R.G. and H.C. (see authors affiliation for address).

**Figure 3.**
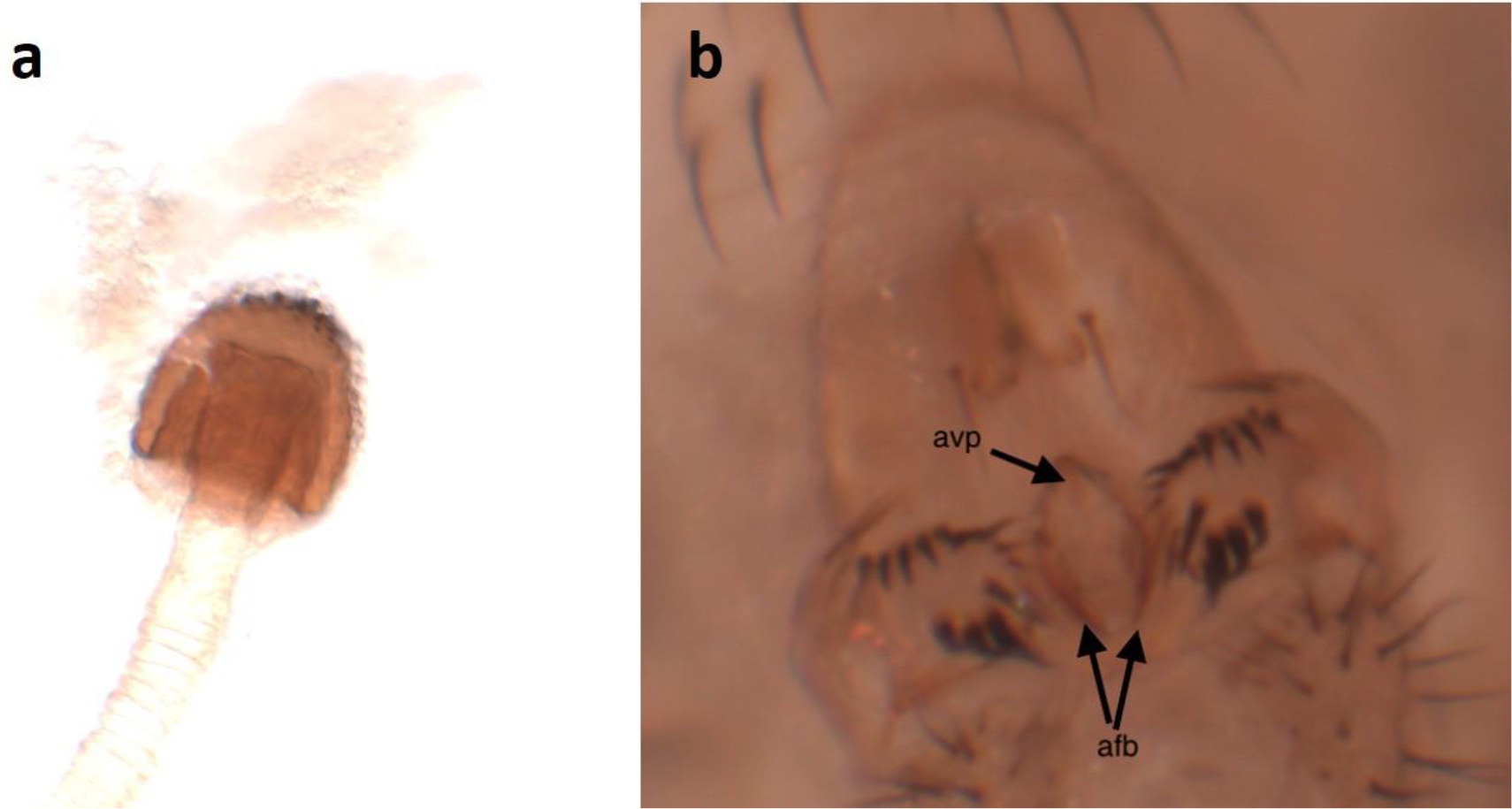
Detail of female spermathecae (a) and male aedeagus (b) of trapped individuals of Z. tuberculatus (avp: aedeagal ventral process, afb: aedeagal flap border).

*Zaprionus tuberculatus* is widespread in Africa and the islands of the Indian Ocean (Madagascar, Comoro, Seychelles, Réunion and Mauritius) and Atlantic Ocean (St Helena and Cap Verde) (14,16–19). At the end of 20^th^ century, *Z. tuberculatus* was recorded in Palearctic region from Canary Islands, Egypt, Cyprus, Malta and Israel (19,20). More recently, *Z. tuberculatus* occurred in Turkey (21), Italy (22), Romania (23), Tunisia (24) and Brazil (25).

The first detection of *Z. tuberculatus* occurred in July 2022 in the hedgerow and captures continued until January 2023 with an interruption in November and December (Table 1). In the fig orchard, we trapped *Z. tuberculatus* in August 2022 and then in October 2022, corresponding respectively to the beginning and the end of fruit harvest. These results suggest that *Z. tuberculatus* could remain in natural woody habitats and colonize orchards when fruits come to maturity. To verifiy whether this observation is the result of a temporary accidental introduction or a permanent population, we harvested around 100 ripe and 100 overripe figs from the fig orchard in the end of September 2023. Over 20 male and female individuals emerged from the overripe figs during the two weeks following the harvest confirming the establishment of a permanent population. Our trapping results show a longer period of occurrence than other studies of *Z. tuberculatus* in newly colonized countries. In Turkey, the first records of *Z. tuberculatus* were reported to be in August (21,26), whereas Raspi *et al*. (22) in Italy and Constantina *et al*. (27) in Romania reported that *Z. tuberculatus* flies were found in the traps only during autumn from September to October. To our knowledge, until now the latest captures were reported in mid-December in Turkey (26), so this is the first time that adults of *Z. tuberculatus* are trapped in winter in Europe. During December, mean daily temperature recorded in the surveyed hedgerow was 8.1°C with a minimum of 3°C (Table 1). Several studies showed evidence of climatic niche shift in exotic species during their expansion (28–31). Such niche shift was also demonstrated in other invasive Drosophilidae, *D. suzukii* (32) and *Zaprionus indianus* (33). Our results suggest that *Z. tuberculatus* could experience a climatic niche shift to accommodate to climate colder than in its tropical origin, as also suggested by Cavalcanti *et al*. (25). Moreover, with climate projection indicating that global warming will result in a progression of Mediterranean climate to the north of France during the next decades (34), climatic conditions will become more suitable to a possible northward expansion of *Z. tuberculatus*.

The pest status of *Z. tuberculatus* and its potential risk for agriculture is not clear. In its native range, *Z. tuberculatus* develops on decaying fruits, particularly on fallen rotting figs (35) and was thus not considered as a pest. However, the spread of this species, the fact that it has successfully been reared on 49 different species of tropical fruits (36) should be a point of vigilance. This is particularly important in view of the first emergences we have observed from overripe, but not rotting, figs. Moreover, Balmes and Mouttet (37) reported emergences of *Z. tuberculatus* from imported fruits of *Citrus sinensis* from South Africa and *Litchi sinensis* from Reunion.

*Zaprionus indianus* is an invasive species that is closely related to *Z. tuberculatus* (14). The pest behavior of *Z. indianus* should also be a warning point for the entry of *Z. tuberculatus* into the EU and France, as they may have similar nutritional ecology and invasion routes. In Brazil, *Z. indianus* was reported attacking figs and was responsible for 50% of fig losses of fruits that were still on trees (38,39). The entry in Central and South America and the damages to fig orchards (40,41) led to its inscription on the alert list of the European and Mediterranean Plant Protection Organization (EPPO) in 2016. Under laboratory conditions, Bernardi et al. (42) observed that *Z. indianus* could oviposit in ripe strawberries and the larvae were then able to develop in the berries. Yet, the ability of *Z. indianus* to oviposit and generate offspring in healthy strawberry fruit was clearly facilitated by injuries caused by *D. suzukii* or by mechanical injuries (42). Although no specific report of damage by *Z. indianus* in the EU has been reported so far, considering the damage caused to figs in South America, *Z. indianus* establishment and spread was considered as a threat with possible large economic consequences in Europe (43). Consequently, in 2022, *Z. indianus* was considered by European Food Safety Authority as a potential Union quarantine pest (43). Our results suggest that a particular attention should be paid also on *Z. tuberculatus*. Indeed, we detected males and females *Z. tuberculatus* in a fig orchard during the harvest period where fruits are systematically removed by the producer before rotting as a prophylactic measure against *D. suzukii* (pers. com. from producer). This may suggest a potential ability to develop in healthy fruits. Moreover, climatic conditions in many EU member states and host plant availability in those areas are conducive for establishment of both *Z. indianus* and *Z. tuberculatus*.

## 4. Conclusions

Our study highlights the risks associated with the discovery of this new species and the potential economic consequences on fruit production. We are now breeding the individuals that have recently emerged from the figs under secure laboratory conditions and we will conduct more investigations to know more about the ecology and biology of *Z. tuberculatus* and to clarify if this species can infest healthy fruits, particularly figs, citrus and strawberries.

## Author Contribution Statement

RG and HC conceived and designed research. RG and HC supervised the data collection. RG and AY managed the identification of specimens. RG and HC drafted the manuscript and all authors read, reviewed and approved the manuscript.

## Acknowledgements

We thank the fig producer for the help in collecting the traps during the monitoring and Jeanne Begouen-Demeaux and Damien Gourmelon for their help in the insects sorting. We also thank the AVFF (Association pour la Valorisation de la Framboise Française) for the constitution of the network of producers involved in the monitoring.

